# Pharmacological Modulation of MC4R in the Periaqueductal Gray Does Not Alter Social Behavior

**DOI:** 10.1101/2025.09.17.676826

**Authors:** Natalia Ruan, Alexandra J. Ng, John P. Christianson

## Abstract

Competing motivational drives are integral to survival and encompass a spectrum of internal physiological needs such as hunger to external goals like social connection. Elucidating how these motivational states interact at the neural level is critical to our understanding of adaptive behavior. Hunger-driven behaviors are primarily initiated by agouti-related peptide (AgRP), an inverse agonist of the melanocortin 4 receptor (MC4R), which acts to promote feeding by suppressing MC4R signaling. MC4R is present in the periaqueductal gray (PAG), a midbrain region involved in pain, appetite, and social behavior. A pilot study found a dense concentration of AgRP fibers in the PAG of male rats, suggesting a potential link between hunger-related signaling and social behavior in this region. To investigate the role of AgRP and MC4R in the PAG on social interaction, we conducted a within-subject dose-response experiment using male rats. Subjects received bilateral microinjections (0.5 µL/side) of either vehicle, an MC4R agonist (THIQ), or an antagonist (HS014) into the PAG prior to a five-minute social exploration (SE) test with a novel juvenile conspecific. Contrary to our hypothesis, pharmacological manipulation of MC4R activity in the PAG did not produce significant changes in social interaction time, suggesting that MC4R in the PAG may not directly play a major role in modulating social behavior in rodents. These results highlight the complexity of the neural circuitry involved in social motivation, hunger, and other competing motivational states.

## I. Introduction

Competing motivational drives are present in the everyday lives of humans and most animals. These range from physiological needs, such as hunger, pain reduction, and reproduction, to external needs such as social interaction and achievement. Physiological needs function to sustain a stable internal state with the goal of maintaining the body’s homeostasis (Berridge, 2004). However, given this array of potential competing motivations, it is possible for these needs to be at odds with each other, playing tug of war until one wins. For example, the desire to eat food when hungry may be suppressed by nearby danger, causing the organism to prioritize safety over satiety (Burnett et al., 2016). Motivational states therefore compete for dominance within an organism, ultimately driving behavior and decision making (Burnett et al., 2016). Internal need states are primarily controlled by populations of neurons in the hypothalamus that orchestrate the physiological and behavioral response to thirst, hunger, and arousal (Graebner et al., 2015). The downstream projections of the hypothalamic neurons that regulate hunger signals have shown to be highly influential in other brain regions and behavior (Deem et al., 2022).

The neural circuitry involved in eating is largely governed by agouti-related peptide (AgRP) neurons in the arcuate nucleus of the hypothalamus. The activation of AgRP neurons stimulates an orexigenic, or appetite stimulant, pathway while the activation of proopiomelanocortin (POMC) neurons in the arcuate nucleus stimulate an anorexigenic, or loss of appetite, pathway (Han et al., 2018). AgRP neurons are hormonally activated by ghrelin, which has more receptors on AgRP neurons than POMC neurons. Ghrelin increases feeding by causing the release of neuropeptide Y (NPY) from AgRP neurons, therefore inhibiting POMC neurons (Han et al., 2018). Leptin, secreted by adipose tissue, depolarizes POMC neurons, inhibiting feeding and the activity of AgRP neurons. Inactivation of leptin has been found to cause obesity, demonstrating the profound effects this hormone can have on eating behaviors (Han et al., 2018). Insulin, produced as a result of increased glucose levels in the blood, has further shown to decrease food intake, potentially stimulating POMC neurons.

The activation of AgRP neurons releases AgRP, which acts as an inverse agonist to the postsynaptic melanocortin receptor, MC4R, in neurons in the paraventricular hypothalamic nucleus (PVN) (Han et al., 2018). These neurons induce satiety, and are agonized by alpha-melanocyte-stimulating hormone (α-MSH) released from POMC neurons. AgRP reduces the basal activity of MC4R, opposing the satiety-promoting effects of α-MSH. Activation of MC4R neurons in the PVN suppresses appetite by directly projecting to the lateral parabrachial nucleus (PBN) in the pons of the brain stem. Therefore, the AgRP-PVN circuit is primarily responsible for “the bidirectional control of hunger and satiety” (Han et al., 2018). The PBN further receives excitatory input from the nucleus of the solitary tract (NTS) in the medulla. (Deem et al., 2022).

Through the vagus nerve, the NTS receives signals to induce satiety from enteroendocrine cells such as cholecystokinin (CCK) and serotonin (5-HT) as a result of gut wall stretching (Andermann et al., 2017). AgRP-MC4R interactions therefore promote feeding behavior by blocking these downstream satiety signals, and enhancing hunger cues.

Along with AgRP, MC4R is therefore crucial to regulating appetite and communicating with other brain regions. MC4R and MC4R mRNA expression has been found in a vast range of brain areas in male and female rats, including the periaqueductal gray (PAG) (Gelez et al., 2010, Kishi et al., 2003). Found in the midbrain, the PAG plays a role in pain processing, defensive behaviors, vocalization, and autonomic regulation (Behbehani, 1995). Rodents display avoidance and freezing behaviors as a result of dorsolateral PAG stimulation (Back & Carobrez, 2018). Additionally, the PAG receives ascending pain transmission from spinothalamic fibers, which is then relayed to the medial thalamus and higher cortical structures (Benarroch, 2008). The PAG is further known for its role in descending pain inhibition and has been found to produce analgesia when stimulated as it is highly involved with the body’s endogenous opioid system (Behbehani, 1995). The PAG is a critical brain region in both ascending pain transmission and descending pain modulation.

The PAG’s role in pain modulation is partially mediated by MC4R. Opioidergic receptors are co-expressed with MC4R in the PAG, demonstrating the relevance of melanocortin receptors in pain transmission and perception (Li et al., 2017, Song et al., 2015). Furthermore, administration of a MC4R antagonist, HS014, to the PAG delays pain facilitation and inflammation in rodents experiencing a chronic pain condition (Chu et al., 2012). Multiple studies have shown that the administration of MC4R antagonists alleviates allodynic behavior in rats experiencing neuropathic pain, whereas MC4R agonists, including α-MSH, heightens nociception (Bellasio et al., 2003, Sandman & Kastin, 1981).

Given the multiple functions of MC4R including those related to eating behaviors and pain processing, it is reasonable to explore how these functions come together in the PAG and their impact on behavior. The PAG receives its largest number of afferents from the hypothalamus, with hypothalamic nuclei terminating in different subregions of the PAG (Behbehani, 1995). In a pilot study, we conducted an immunohistochemistry to stain for AgRP, the inverse agonist of MC4R, in rat brain tissue (Figure 1). A high concentration of AgRP fibers was found in the PAG, guiding the direction of this work to investigate the role of MC4R in rodent behavior, specifically in social interaction. It is known that MC4R agonists facilitate the formation of partner preferences in male and female prairie voles, however it is unknown whether MC4R plays a role in basic social interaction (Modi et al., 2015). To this end, we investigated to what extent antagonizing and agonizing the MC4R in the PAG impacted sociability in rats using a social exploration test.

**Figure 1:**
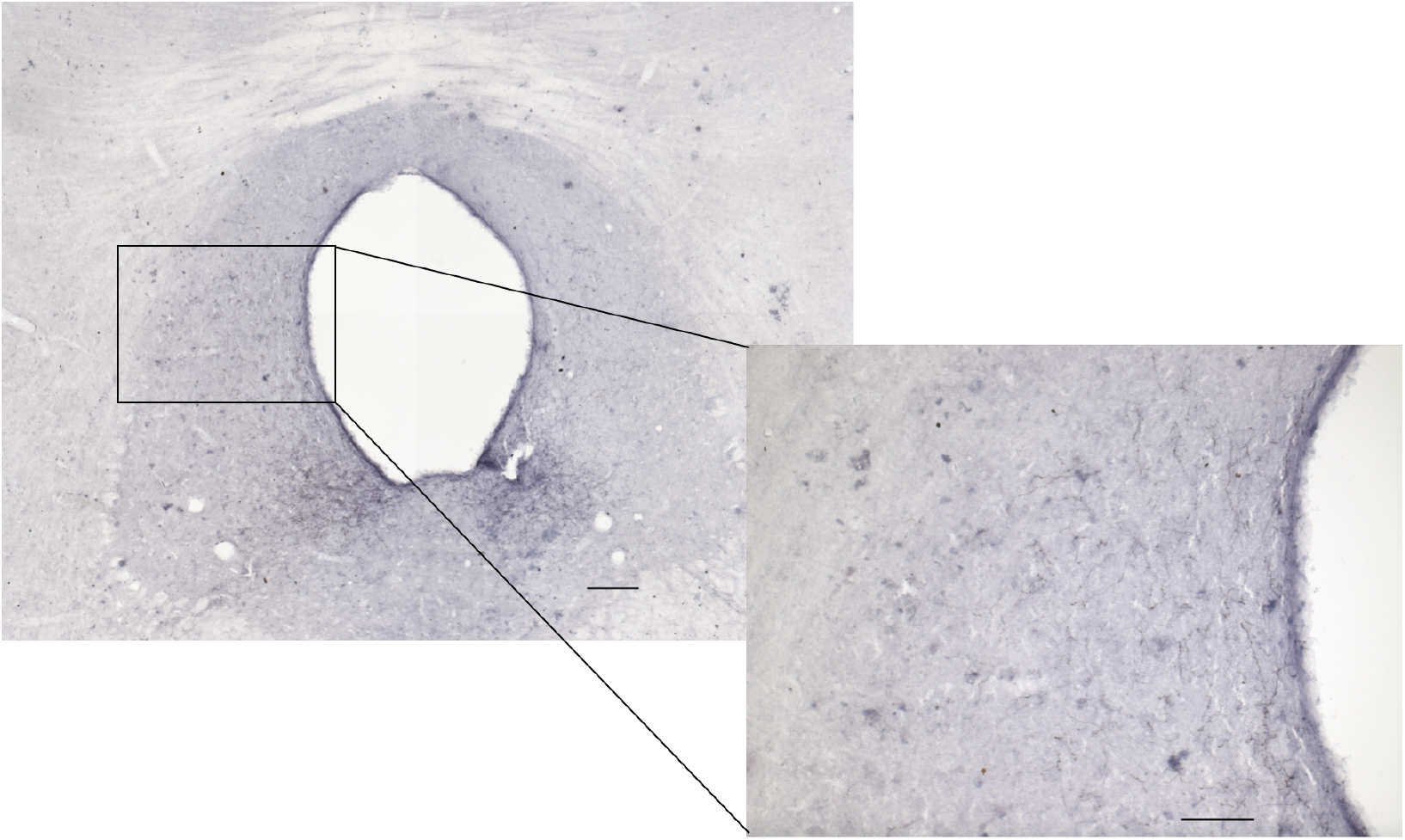
Immunohistochemical staining of AgRP fibers in the periaqueductal gray using a human anti-AgRP antibody. Scale bar = 200µm for larger field of view. Scale bar = 100µm for smaller field of view.

**Figure 2:**
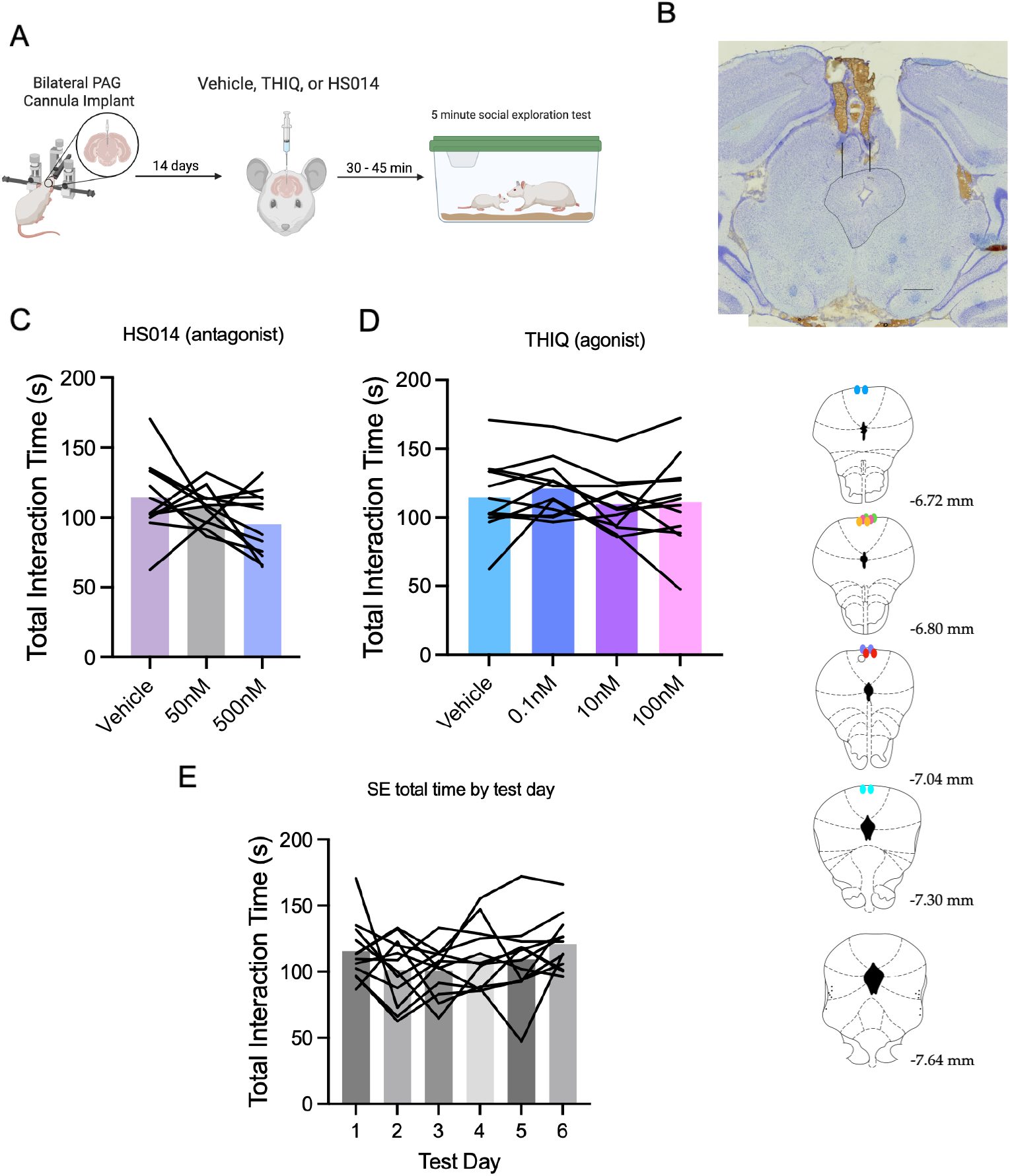
MC4R receptors in the PAG are not necessary for social behavior. **A**. Schematic representation of the experimental protocol. Bilateral cannula were implanted in the periaqueductal gray and rats recovered for two weeks. Following recovery, rats received a bilateral microinjection (0.5µL/side) of vehicle or THIQ/HS014 and underwent a social exploration test with a naive male juvenile conspecific 30-45 minutes after injections. **B**. Image of the bilateral cannula tract on top of the periaqueductal gray, outlined in the black circle. The full reach of the injectors is represented by the black lines. Below, the maps of the cannula placements of the PAG microinjections are represented. Each color represents each test rat (n = 7). **C**. Average interaction time (s) of test rats during the first three days of testing following microinjection of vehicle or HS014 (50 nM, 500 nM). Each line represents the social interaction time of individual test rats. No significant difference was found between vehicle and antagonist conditions (F_TREATMENT_ (1.996,11.98) = 0.9270, P = 0.4222). **D**. Average interaction time (s) of test rats in the last three days of testing following microinjection of vehicle or THIQ (0.1 nM, 10 nM, 100 nM). Administration of THIQ or vehicle caused no significant difference in interaction time (F_TREATMENT_ (2.196, 13.17) = 1.025, P = 0.3923). **E**. Average interaction time (s) of each rat during the social exploration (SE) test, represented by individual lines, across each test day. No significant difference was found in interaction time across testing days (F_TEST DAY_ (3.091, 18.55) = 1.326, P = 0.2961).

## II. Methods

### Animals

Male Sprague Dawley rats were obtained from Charles River Laboratories at 244-250g. Once they arrived at the Boston College Animal Care Facility, they acclimated for one week in the vivarium and were housed in pairs. The animals were given food and water ad libitum and were maintained on a 12-h light/dark cycle. All behavioral testing occurred in the first four hours of their light cycle. The Boston College Institution Animal Care and Use Committee approved of the following procedures, which were carried out in accordance with the National Institute of Health’s *Guide for the Care and Use of Laboratory Animals*.

### Surgery

Bilateral cannula were surgically inserted in the PAG (from bregma: A/P −7.5 mm, M/L +/-0.5 mm, D/V: −5 mm) in a stereotaxic frame while rats were under isoflurane anesthesia (1.5-5% v/v in O_2_, Somnoflo). Stainless steel screws and acrylic dental cement were used to secure the cannula. Each rat was injected with meloxicam (1 mg/kg), lactated Ringer’s solution, and antibiotic penicillin following the surgical procedure. On the days following surgery, rats were given 0.06 mL of meloxicam as an analgesic, and weights were continuously recorded to ensure their physical health. If the rats had lost more than 6 grams of weight, or 2% of their pre-surgery body weight, 5 mL of lactated Ringer’s solution was administered. Rats recovered for 2 weeks before behavioral testing in their home cages. A week prior to behavioral testing, test animals were habituated to being wrapped in a cloth towel in order to simulate cannula injections and reduce stress associated with the experimental protocol.

### Pharmacological manipulations

The MC4R antagonist, HS014, and the agonist, THIQ, were obtained from Tocris Bioscience. HS014 has previously been used in rodents to stimulate feeding behavior through cannula microinjections (Kask & Schiöth, 2000). Multiple studies have further used HS014 as a MC4R antagonist and have found an effect through differing delivery methods (Chu et al., 2012, Bellasio et al., 2003, Shikdar & Alghamdi, 2021). THIQ was chosen as a selective MC4R agonist as it has also been used in multiple studies to activate the MC4R (Ross et al., 2023,

Starnowska-Sokół et al., 2020). 1 mg of HS014 was diluted in 1 mL of deionized water to create a 639.48 µM stock solution. A 500 nM solution of HS014 was created by diluting 7.8 µL of the stock solution into 10 mL of saline. 1.6 µL of the stock solution was diluted into 20 mL of saline to create a 50 nM solution of HS014. THIQ doses of 0.1, 10, and 100 nM were created using a 1.6973 mM stock solution of THIQ in 1 mL of DMSO. Smaller doses of the THIQ agonist were chosen as prior studies have indicated highly potent effects with miniscule doses of THIQ (Richardson et al., 2013). Test animals were given saline/DMSO solution as a vehicle. This solution was administered along with varying doses of the agonist and antagonist 30-45 minutes before testing.

### Social Exploration Test

Six days of one-on-one social interaction tests took place. In the first three days, rats received a bilateral microinjection (0.5µL/side) into the PAG of a vehicle or two doses of the antagonist (HS014; 50nM and 500nM) directly into the PAG in a counterbalanced manner. Five minutes of social interaction with a naive juvenile male rat occurred 30-45 minutes after microinjections. Two trained observers recorded the total interaction time of the test rat in seconds. Social interaction was measured as the amount of time the test rat spent interacting with the conspecific. This included direct sniffing, touching, and/or licking the conspecific. All social exploration tests were recorded on video. The next three days followed the same protocol using microinjections of vehicle and the agonist at multiple doses (0.1nM, 10 nM and 100nM) in a counterbalanced manner. Increasing doses of the antagonist and agonist allowed for a within-subject dose response experiment.

### Tissue Collection and Slicing

Three days following testing, rats were overdosed with tribromoethanol (Sigma), brains were extracted, flash frozen, and sliced in 40 µm sections using a freezing cryostat (Leica). Slices were immediately mounted on gelatin-coated slides (Fisher), cresyl-violet stained, and coverslipped with Permount. Cannula placements were verified using the rat whole brain stereotaxic atlas as a reference (Paxinos & Watson, 1998) when visualizing microinjector tip locations on the microscope. Rats with absent or incorrect cannula locations (Bregma −6.7 through Bregma −8.2) were excluded from subsequent statistical analyses.

### Statistical Analysis

Interaction times during the social exploration test were analyzed using a one-way analysis of variance (ANOVA) with Tukey’s multiple comparisons test. Differences in social interaction depending on agonist/antagonist dose were considered significant at p < 0.05.

## III. Results

### Antagonizing and agonizing MC4R in the PAG does not impact total social interaction time with a naive juvenile conspecific

In order to determine if MC4R in the PAG play a role in social behavior in rodents, rats underwent a dose response experiment investigating whether varying doses of the MC4R antagonist (HS014) and agonist (THIQ) impacted social interaction time during a five minute social exploration test. A one-way ANOVA demonstrated no significant main effect between the administration of vehicle and increasing doses of HS014 in total interaction time with a naive juvenile conspecific (F_TREATMENT_ (1.996,11.98) = 0.9270, P = 0.4222). Additionally, individual differences between subjects were not significant (F_SUBJECT_ (6, 12) = 1.942, P = 0.1544). While increasing doses of HS014 demonstrated a downward trend in total interaction time, suggesting the antagonist’s influence on decreasing sociability, these results were not significant. Although the one-way ANOVA did not reveal statistically significant differences, analyses of pairwise comparisons were conducted in an exploratory manner to assess potential trends between the treatment groups for the administration of HS014. Pairwise comparisons indicated no significant differences between any of the treatment groups (vehicle vs 50nM: P = 0.9909; vehicle vs 500nM: P = 0.5366; 50nM vs 500nM: P = 0.4890).

Additionally, a separate one-way ANOVA showed that increasing doses of THIQ did not have a significant main effect on social interaction time in comparison with vehicle (F_TREATMENT_ (2.196, 13.17) = 1.025, P = 0.3923). There was no significant effect between subjects (F_SUBJECT_(6, 18) = 2.747, P = 0.0449). Subsequent analysis of pairwise comparisons demonstrated no significant difference between treatment groups (vehicle vs 0.1nM: P = 0.4427; vehicle vs 10nM: P = 0.9970; vehicle vs 100nM: P = 0.9660; 0.1nM vs 10 nM: P = 0.4823; 0.1nM vs 100nM: P = 0.5920; 10nM vs 100nM: P = 0.9865).

To assess if there were significant differences in total interaction time across each test day, a one-way ANOVA was run. No significant difference was found (F_TEST DAY_ (3.091, 18.55) = 1.326, P = 0.2961), suggesting that repeated social interactions with the naive juvenile conspecifics over the six test days or order of drug administration did not impact the experimental results.

## IV. Discussion

The present study examined whether antagonism or agonism of the MC4R in the PAG influences social behavior in rodents. Given the presence of AgRP fibers in the PAG as well as previous research demonstrating the prosocial effects of MC4R agonism (Modi et al., 2015), we hypothesized that the administration of the MC4R agonist, THIQ, would increase interaction time in rats during a social exploration test. The administration of HS014 as an antagonist to the MC4R was expected to either decrease or have no effect on interaction time. Contrary to this hypothesis, no significant differences in interaction time with a naive juvenile conspecific were observed across increasing doses of THIQ (0.1 nM, 10 nM, 100 nM) and HS014 (50 nM, 500 nM) compared to vehicle. This suggests that the MC4R in the PAG does not play a crucial role in social behavior in rodents.

However, the present study utilized two drugs at a limited range of doses, leaving open the possibility that an alternative, optimal dose may have yielded the desired effect. It is also possible that choosing an alternative agonist/antagonist with more selective properties could have changed the results. Nevertheless, the decision to use HS014 as a MC4R antagonist and THIQ as a MC4R agonist was supported by the literature, as these compounds have been consistently used to elicit specific behavioral responses in rodents. Furthermore, the immunohistochemistry conducted in the pilot study displayed dense AgRP fiber labeling in the ventral PAG (Figure 1), particularly around the aqueduct. The cannula tips targeted the dorsal region of the PAG for drug injections. While it is likely that the drugs diffused far from the injection site, it is essential to take this into account when interpreting the results of this study and suggesting future improvements.

Given that melanocortin receptors are involved in feeding, pain processing, and social behavior, the results of this study call into question other avenues that can be explored involving the receptor’s role in the PAG. Much research has focused on the significance of the MC4R within the physiology of pain (Behbehani, 1995; Benarroch, 2008; Pagano et al., 2012; Ye et al., 2014). Past findings demonstrate the critical role of MC4R signaling in the PAG as well as the effects of MC4R agonism and antagonism on pain perception (Bellasio et al., 2003; Chu et al., 2012; Sandman & Kastin, 1981). Future studies could use a somewhat similar paradigm to the present study, examining how modulating pain perception through MC4R agonism and antagonism changes social behavior in rodents. This approach would provide deeper insight into the role of the PAG in balancing competing motivational drives.

MC4R and AgRP are further involved in feeding behavior, an area that warrants continued investigation. Beyond the PAG, the function of MC4R throughout specific brain regions has been explored in relation to internal need states such as hunger. Past research has shown that MC4Rs in the PVN and potentially in the amygdala contribute to food intake, while separate MC4Rs in other brain areas still unknown play a role in energy expenditure (Balthasar et al., 2005; Chen et al., 2000). The region specificity of MC4R function can guide future work assessing how the feeling of hunger and/or food intake interacts with a competing motivational drive such as social interaction. Similarly, investigating other neural circuits involving AgRP could elucidate the underlying mechanisms involved in hunger and feeding, as it has been shown that AgRP is necessary for these processes (Krashes et al., 2011). AgRP-expressing neurons communicate with reward centers in the brain and inhibit regions involved in energy-demanding processes such as the stress response and reproduction (Han et al., 2018). It is evident that AgRP plays a role in diverting an organism’s resources towards food seeking behavior, therefore future research is necessary to understand how this internal drive interacts with competing motivational states.

Investigating the role of AgRP and the function of MC4R has potential therapeutic implications. In regards to pain physiology, HS014 has previously been used as an MC4R antagonist to alleviate neuropathic pain in a rodent model (Korczeniewska et al., 2021). Although much remains unknown, further investigation into the neural circuitry of the PAG and the effects of pharmacological manipulations could provide deeper insight into the role of MC4R as a therapeutic target in pathological pain. Additionally, continuing to explore the brain regions and neural circuits in which AgRP exerts its effects could shed light on how deficits in this circuitry could underlie psychopathologies such as anorexia nervosa and binge eating disorder. The overactivation or underactivation of the neural circuitry involving AgRP could lead to hyperphagia and appetite suppression, respectively (Han et al., 2018). Future studies exploring how alterations in AgRP function across different brain regions influences feeding behavior and motivational drives in rodents could enhance our understanding of the neural basis of eating-related psychopathologies.

The neural circuitry involving AgRP and the function of its receptor, MC4R, play a role in essential physiological processes including feeding behavior and pain perception. Continuing to explore how these processes interact with other, non-physiological drives such as social interaction could provide deeper insight into the complex regulation of behavior. The present study serves as a valuable starting point in the investigation of how competing motivational drives integrate to produce behavioral responses.

## Notes

### Competing Interest Statement

The authors have declared no competing interest.

### Funding

- National Institutes of Health: MH119422, MH133304

